# Scientometric Trends for Coronaviruses and Other Emerging Viral Infections

**DOI:** 10.1101/2020.03.17.995795

**Authors:** Dima Kagan, Jacob Moran-Gilad, Michael Fire

**Affiliations:** Department of Software and Information Systems Engineering, Ben-Gurion University of the Negev; Department of Health Systems Management, Faculty of Health Sciences, Ben-Gurion University of the Negev

**Keywords:** Coronavirus, Emerging viruses, Epidemics, SARS

## Abstract

COVID-19 is the most rapidly expanding coronavirus outbreak in the past two decades. To provide a swift response to a novel outbreak, prior knowledge from similar outbreaks is essential. Here, we study the volume of research conducted on previous coronavirus outbreaks, specifically SARS and MERS, relative to other infectious diseases by analyzing over 35 million papers from the last 20 years. Our results demonstrate that previous coronavirus outbreaks have been understudied compared to other viruses. We also show that the research volume of emerging infectious diseases is very high after an outbreak and drops drastically upon the containment of the disease. This can yield inadequate research and limited investment in gaining a full understanding of novel coronavirus management and prevention. Independent of the outcome of the current COVID-19 outbreak, we believe that measures should be taken to encourage sustained research in the field.

## 1 Introduction

Infectious diseases remain a major cause of morbidity and mortality worldwide, in developed countries and particularly in the developing world [1]. According to the World Health Organization, out of the top-10 causes of death globally, three are infectious diseases [1]. In light of the continuous emergence of infections, the burden of infectious diseases is expected to become even greater in the near future [2, 3]. Many emerging pathogens are RNA viruses, and notable examples over the last two decades include the SARS coronavirus in 2002-2003 in China, pandemic influenza (swine flu) A/H1N1 in 2009, the MERS coronavirus in 2012 in the Middle East, and Ebola virus disease in 2013-2014 in Africa.

Currently, the world is struggling with a novel strain of coronavirus (SARS-CoV-2) that emerged in China during late 2019 and by the time of this writing has infected more than 4,400,000 people and killed more than 302,000 [4, 5]. COVID-19 is the latest and third serious human coronavirus outbreak in the past 20 years. Additionally, of course, there are several more typical circulating seasonal human coronaviruses causing respiratory infections. It is still too early to predict the epidemic course of COVID-19, but it is already a pandemic that appears more difficult to contain than its close relative SARS-CoV [6, 7].

Much can be learned from past infectious disease outbreaks to improve preparedness and response to future public health threats. Three key questions arise in light of the COVID-19 outbreak: *To what extent were the previous human coronaviruse (SARS and MERS) outbreaks studied? Is research on emerging viruses being sustained, aiming to understand and prevent future epidemics? Are there lessons from academic publications on previous emerging viruses that could be applied to the current COVID-19 epidemic?*

In this study, we answer these vital questions by utilizing state-of-the-art data science tools to perform a large-scale analysis of 35 million papers, of which 1,908,211 concern the field of virology. We explore nearly two decades of infectious disease research published from 2002 up to today. We particularly focus on public health crises, such as SARS, influenza (including seasonal, pandemic H1N1, and avian influenza), MERS, and Ebola virus disease, and compare them to HIV/AIDS and viral hepatitis B and C, three bloodborne viruses that are associated with a significant global health burden for more than two decades.

A crucial aspect of being prepared for future epidemics is sustained ongoing research of emerging infectious diseases even at ‘times of peace’ when such viruses do not pose an active threat. Our results demonstrate that research on previous coronaviruses, such as SARS and MERS, was conducted by a relatively small number of researchers centered in a small number of countries, suggesting that such research could be better encouraged. We propose that regardless of the fate of COVID-19 in the near future, sustained research efforts should be encouraged to be better prepared for the next outbreak.

## 2 Background

This research is a large-scale scientometric study in the field of infectious diseases. We focus on the quantitative features and characteristics of infectious disease research over the past two decades. In this section, we present studies that analyze and survey real-world trends in the field of infectious diseases (see the Infectious Disease Trends subsection) and studies that relate to bibliometric trends in general and public health in particular (see the Bibliometric Trends subsection).

### 2.1 Infectious Disease Trends

There is great promise in utilizing big data to study epidemiology [8]. One approach is to gather data using different surveillance systems. For example, one such system is ProMED. ProMED was launched 25 years ago as an email service to identify unusual worldwide health events related to emerging and reemerging infectious diseases [9]. It is used daily around the globe by public health policy makers, physicians, veterinarians, and other healthcare workers, researchers, private companies, journalists, and the general public. Reports are produced and commentary is provided by a global team of subject-matter experts in a variety of fields. ProMED has over 80,000 subscribers and over 60,000 cumulative event reports from almost every country in the world. Additionally, there are many different systems used by different countries and health organizations worldwide.

In 2006, Cowen et al. [10] evaluated the ProMED dataset from the years 1996 to 2004. They discovered that there are diseases that received more extensive coverage than others; “86 disease subjects had thread lengths of at least 10 reports, and 24 had 20 or more.” They note that the pattern of occurrence is hard to explain even by an expert in epidemiology. Also, with the level of granularity of ProMED data, it is very challenging to predict the frequency that diseases are going to accrue. In 2008, Jones et al. [2] analyzed the global temporal and spatial patterns of emerging infectious diseases (EIDs). They analyzed 305 EIDs between 1940 and 2004 and demonstrated that the threat of EIDs to global health is increasing. The same year, Freifeld et al. [11] developed HealthMap, an interactive surveillance system that integrates disease outbreak reports from various sources.

Data about infectious diseases can also come from web- and social-based sources. For instance, in 2009, Ginsberg et al. [12] used Google search queries to monitor the spread of influenza epidemics. They used the fact that many people search online before going to doctors, and they found that during a pandemic, the volume of searches differs from normal. They then created a mathematical model to forecast the spread of flu. This research was later converted into a tool called Google Flu Trends, and at its peak, Google Flu Trends was deployed in 29 countries worldwide. However, not everything worked well for Google Flu Trends; in 2009, it underestimated the flu volume, and in 2013, it predicted more than double the number of cases than the true volume [13]. As a result of such discrepancies, Google shut down the Google Flu Trends website in 2015 and transferred its data to academic researchers [14]. Also in 2009, Carneiro and Mylonakis [15] used large amounts of data to predict flu outbreaks a week earlier than prevention surveillance systems.

In 2010, Lampos and Cristianini [16] extended the idea of Carneiro and Mylonakis [15] to use temporal data to monitor outbreaks. Instead of using Google Trends, they used Twitter as their data source. They collected 160,000 tweets from the UK, and as ground truth, they used HPA weekly reports about the H1N1 epidemic. Using textual markers to measure flu on Twitter, they demonstrated that Twitter can be used to study disease outbreaks, similar to Google Trends. Also the same year, Salathé and Khandelwal [17] analyzed Twitter and demonstrated that it is possible to use social networks to study not only the spread of infectious disease but also vaccinations. They found a correlation between the sentiment in tweets toward an influenza vaccine and the vaccination rate.

In 2014, Generous et al. [18] used Wikipedia to monitor and forecast infectious disease outbreaks. They examined Wikipedia access logs to forecast outbreak volumes for 14 combinations of diseases and locations. The model worked successfully for only 8 out of the 14 cases. Also, the authors suggested that it was even possible to transfer a model between locations without retraining it. In contrast to most of the web-based disease monitoring methods, Wikipedia-based monitoring presents a fully open forecasting system that can be easily reproducible. Generally, in the past couple of years, Wikipedia has become a widely used data source for medical studies [19, 20]. Moreover, a recent report [21] shows that Wikipedia has successfully kept itself clean from the misinformation spread during the COVID-19 outbreak. In 2015, Santillana et al. [22] took the influenza surveillance one step further by fusing multiple data sources. They used five datasets: Twitter, Google Trends, near real-time hospital visit records, FluNearYou, and Google Flu Trends. They used all these data sources with a machine-learning algorithm to predict influenza outbreaks. In 2017, McGough et al. [23] dealt with the problem of significant delays in the publication of official government reports about Zika cases. To solve this problem, they used the combined data of Google Trends, Twitter, and the HealthMap surveillance system to predict estimates of Zika cases in Latin America.

In 2018, Breugelmans et al. [24] explored the effects of publishing in open access journals and collaboration between European and sub-Saharan African researchers in the study of poverty-related disease. To this end they used the PubMed dataset but discovered it is not suited to performing full bibliometric analysis; to deal with this issue they also utilized Web Of Science as a data source. They discovered that there is an advantage for open access publications in terms of citations. In 2020, Head et al. [25] studied infectious disease funding. They discovered that HIV/AIDS is the most funded disease. Additionally, they discovered a pattern where Ebola, Zika, influenza, and coronavirus funding were highest after an outbreak.

There is substantial controversy surrounding the use of web-based data to predict the volume of outbreaks. The limitations of Google Flu Trends, mentioned above, raised the question of reliability of social data for assessing disease spread. Lazer [26] noted that these types of methods are problematic since companies like Google, Facebook, and Twitter are constantly changing their products. Studies based on such data sources may be valid today but not be valid tomorrow, and may even be unreproducible.

### 2.2 Bibliometric Trends

In 2005, Vergidis et al. [27] used PubMed and JCR (Journal Citation Reports) to study trends in microbiology publications. They discovered that microbiology research in the US had the highest average impact factor, but in terms of research production, Western Europe was first. In 2008, Uthman [28] analyzed trends in paper publications about HIV in Nigeria. He found growth (from 1 to 33) of the number of publications about HIV in Nigeria and that papers with international collaborations were published in journals with a higher impact factor. In 2009, Ramos et al. [29] used Web of Science to study publications about infectious diseases in European countries. They found that more papers in total were published about infectious diseases in Europe than in the US.

In 2012, Takahashi-Omoe and Omoe [30] surveyed publications of 100 journals about infectious diseases. They discovered that the US and the UK had the highest number of publications, and relative to the country’s socioeconomic status, the Netherlands, India, and China had relatively high productivity. In 2014, similar to Wislar et al. [31], Kennedy et al. [32] studied ghost authorship in nursing journals instead of biomedical journals. They found that there were 27.6% and 42% of ghost and honorary authorships, respectively.

In 2015, Wiethoelter et al. [33] explored worldwide infectious disease trends at the wildlife-livestock interface. They found that 7 out of the top 10 most popular diseases were zoonoses. In 2017, Dong et al. [34] studied the evolution of scientific publications by analyzing 89 million papers from the Microsoft Academic dataset. Similar to the increase found by Aboukhalil [35], they also found a drastic increase in the number of authors per paper. In 2019, Fire and Guestrin [36] studied the over-optimization in academic publications. They found that the number of publications has ceased to be a good metric for academic success as a result of longer author lists, shorter papers, and surging publication numbers. Citation-based metrics, such as citation number and h-index, are likewise affected by the flood of papers, self-citations, and lengthy reference lists.

### 3 Data Description

In this study, we fused four data sources to extract insights about research on emerging viruses. In the rest of this subsection we describe these data sources.

1. **MAG** - Microsoft Academic Graph is a dataset containing “scientific publication records, citation relationships between those publications, as well as authors, institutions, journals, conferences, and fields of study” [37]. The MAG dataset we used was from 22 March 2019 and contains data on over 210 million papers [38]. This dataset was used as the main dataset of the study. Similar to Fire and Guestrin [36], we only used papers that had at least 5 references in order to filter non peer-reviewed publications, such as news columns which are published in journals.
2. **PubMed** - PubMed is a dataset based on the PubMed search engine of academic publications on the topics of medicine, nursing, dentistry, veterinary medicine, health care systems, and preclinical sciences [39]. One of the major advantages of using the PubMed dataset is that it contains only medical-related publications. The data on each PubMed paper contains information about its venue, authors, and affiliations, but it does not contain citation data. In this study, we used the 2018 annual baseline PubMed dataset containing 29,138,919 records[40]. We mainly utilized the PubMed dataset to analyze journal publications (see Paper Trends Section).
3. **SJR** - Scientific Journal Rankings is a dataset containing the information and ranking of over 34,100 journals from 1999 to 2018 [41], including their SJR indicator,^1^ the best quartile of the journal,^2^ and more. We utilized the SJR dataset to compare the rankings of different journals to assess the level of their prestige.
4. **Wikidata** - Wikidata is a dataset holding a vast knowledge about the world, containing data on over 78,252,808 items [44]. Wikidata stores metadata about items, and each item has an identifier and can be associated with other items. We utilized the Wikidata dataset to extract geographic information for academic institutions in order to match a paper with its authors’ geographic locations.

## 4 Analyses

### 4.1 Infectious Disease Analysis

To study the research of emerging viruses over time, we analyzed the datasets described in the Data Description section. In pursuing this goal, we used the code framework recently published by Fire and Guestrin [36], which enables the easy extraction of the structured data of papers from the MAG dataset. The MAG and PubMed datasets were filtered according to a predefined list of keywords. The keyword search was performed in the following way: given a set of diseases *D* and a set of papers *P*, from each paper title *p*_*t*_, where *p* ∈ *P*, we created a set of *word-grams. Word-grams* are defined as *n-grams* of words, i.e., all the combinations of a set of words in a phrase, without disrupting the order of the words. For example, the *word-grams* of the string “Information on Swine Flu,” *word*-*grams*(Information on Swine Flu), will return the following set: {*Information, on, Swine, Flu, Information on, on Swine, Swine Flu, Information on Swine, on Swine Flu, Information on Swine Flu*}. Next, for each *p*, we calculated *word*-*gram*(*p*_*t*_) ∩ *D*, which was considered as the diseases with which the paper was associated.

In the current study, we focused on the past emerging coronaviruses (SARS and MERS). There are many other strains of the human coronavirus, and four of them are known for causing seasonal respiratory infections [45]. We focused on SARS and MERS since they are closer to SARS-CoV-2 and both have zoonotic origins and raised international public health concern. Additionally, we also analyzed Ebola virus disease, influenza (seasonal, avian influenza, swine flu), HIV/AIDS, hepatitis B, and hepatitis C as comparators that represent other important emerging infectious diseases from the past two decades. For these nine diseases, we collected all their aliases, which were added to the set of diseases *D* and were used as keywords to filter the datasets. To reduce the false-positive rate, we analyzed only papers that, according to the MAG dataset, were in the categories of medicine or biology, and following Fire and Guestrin [36] had at least five references. Additionally, to explore the trend in the core categories of infectious disease research, we performed the same analysis on the virology category. In the rest of this section, we describe the specific calculations and analyses we performed.

#### 4.1.1. Paper Trends

To explore the volume of studies on emerging viruses, we examined the publication of papers about infectious diseases. First, we defined several notions that we used to define publication and citation rates. Let *D* be a set of disease names and *P* a set of papers. Namely, for a paper *p* ∈ *P, p*_*Disease*_ is defined as the disease that matches the paper’s keywords, *p*_*year*_ as the paper’s publication year, and *p*_*citations*_ as the set of papers citing *p*. Using these notions, we defined the following features:

- *Number of Citations* - the total number of citations for a specific infectious disease.
- *Number of Papers* - the total number of published papers for a specific infectious disease.
- *Normalized Citation Rate* (*NCR*_*y*_) - the ratio between the *Number of Citations* on a specific infectious disease *d* and the total number of citations about medicine or biology in year *y*.^3^

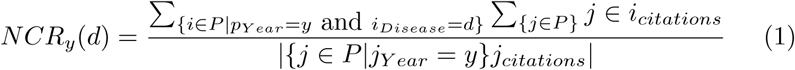
- *Normalized Paper Rate* (NPR) - the ratio between the *Number of Papers* published on a specific infectious disease *d* to the total number of papers in the fields of medicine or biology in the year *y*.

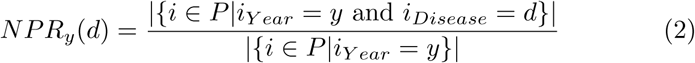

Using these metrics, we inspected how the coronavirus publication and citation rates differed from other examined EIDs. We analyzed how trends of citations and publications have changed over time. Additionally, to inspect the similarities between the trends of different diseases we calculated the DTW (Dynamic time warping) distance [46] between all the disease pairs. Finally, we clustered the time-series using TimeSeriesKMeans [47]

#### 4.1.2 Journal Trends

To investigate the relationship between journals and their publication of papers about emerging viruses, we combined the Semantic Scholar and PubMed datasets with the SJR dataset using ISSN, and selected all the journals from SJR categories related to infectious diseases (immunology, epidemiology, infectious diseases, virology, and microbiology). First, we inspected whether coronavirus papers are published in the top journals. We selected the top-10 journals by SJR and calculated the number of papers they had published for each disease over time. Next, we inspected how published papers about coronavirus are regarded relative to other EIDs in terms of ranking. To this end, we defined a new metric, *JScore*_*t*_. *JScore*_*t*_ is defined as the average SJR score of all published papers on a specific topic *t*. We used *JScore*_*t*_ to observe how the prominence of each disease in the publication world has changed over time. Lastly, we explored publications by looking at the quartile ranking of the journal over time.

#### 4.1.3 Author Trends

To study how scientific authorship has changed in the field of infectious diseases, we explored what characterizes the authors of papers on different diseases. We inspected the number of new authors over time to check how attractive emerging viruses are to new researchers. Additionally, we analyzed the number of experienced authors, where author experience is defined as the time that has passed from his or her first publication. The authors were identified by the identification number provided in the MAG dataset. Author disambiguation is a challenging task; Microsoft combined multiple methods to generate their author identifications [48]. We also analyzed the number of authors who wrote multiple papers about each disease.

#### 4.1.4 Collaboration Trends

To inspect the state of international collaborations in emerging virus research, we mapped academic institutions to geolocation. However, it is not a trivial task to match institution names. Institution names are sometimes written differently; for example, Aalborg University Hospital and Aalborg University are affiliated. However, there are cases where two similar names refer to different institutions; for example, the University of Washington and Washington University are entirely different institutions. To deal with this problem, we used the affiliation table in the MAG dataset. To determine the country and city of each author, we applied a five-step process:

1. For each institution, we looked for the institution’s page on Wikidata. From each Wikidata page, we extracted all geography-related fields.^4^
2. To first merge all the Wikidata location fields, we used the “coordinate location” with reverse geocoding to determine the city and country of the institution.
3. For all the institutions that did not have a “coordinate location” field, we extracted the location data from the other available fields. We crossed the data against city and country lists from GeonamesCache Python library [49] to determine whether the data in the field described a city or a country.
4. To acquire country data for an institution that had only city data on Wikidata, we used GeonamesCache city-to-country mapping lists.
5. To get city and country data for institutions that did not have the relevant fields on Wikidata, we extracted geographic coordinates from Wikipedia.org.^5^ Even though Wikidata and Wikipedia.org are both operated by the Wikimedia Foundation, they are independent projects which have different data. Similar to Wikidata coordinates, we used reverse geocoding to determine the city and country of the institution.

Using the extracted geodata, we explored how international collaborations change over time in coronavirus research. Finally, we explored which countries have the highest number of papers about coronavirus and which countries have the highest number of international collaborations over time.

## 5 Results

In the following subsections, we present all the results of the experiments which were described in the Analyses section.

### 5.1 Results of Paper Trends

In recent years, there has been a surge in academic publications, yielding more than 1 million new papers related to medicine and biology each year (see Figure 1a). In contrast to the overall growth in the number of infectious disease papers, there has been a relative decline in the number of papers about the coronaviruses SARS and MERS (see Figure 1b). Also, we found that 0.4% of virology studies in our corpus from the past 20 years involved human SARS and MERS, while HIV/AIDS accounts for 7.9 % of all virology studies. We observed that, unlike the research in the domain of HIV/AIDS and avian influenza that has been published at a high and steady pace over the last 20 years, SARS was studied at an overwhelming rate after the 2002-2004 outbreak and then sharply dropped after 2005 (Figure 2). In terms of *Normalized Paper Rate* (see Figure 2), after the first SARS outbreak, there was a peak in publishing SARS-related papers with NPR twice as high as Ebola’s. However, the trend dropped very quickly, and a similar phenomenon can be observed for the swine flu pandemic. The MERS outbreak achieved a much lower NPR than SARS, specifically more than 16 times lower when comparing the peaks in SARS and MERS trends. In terms of *Normalized Citation Rate* (Figure 3), we observed the same phenomenon as we did with NPR. Observing Figures 9 and 10, we can see that there are diseases with very similar trends. More precisely, NPR and NCR trends are in two clusters, where the first cluster contains avian influenza, Ebola, MERS, SARS, and swine flu, and the second cluster contains HIV/AIDS, hepatitis B, hepatitis C, and influenza.

**Figure 1:**
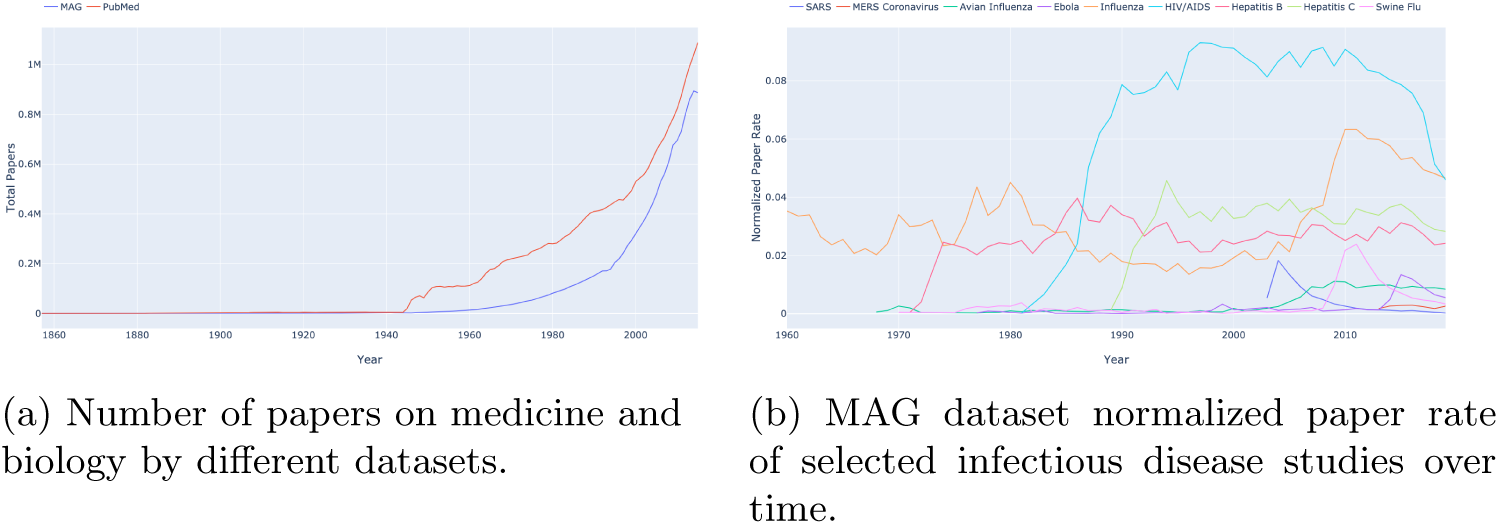
The number of papers over time.

**Figure 2:**
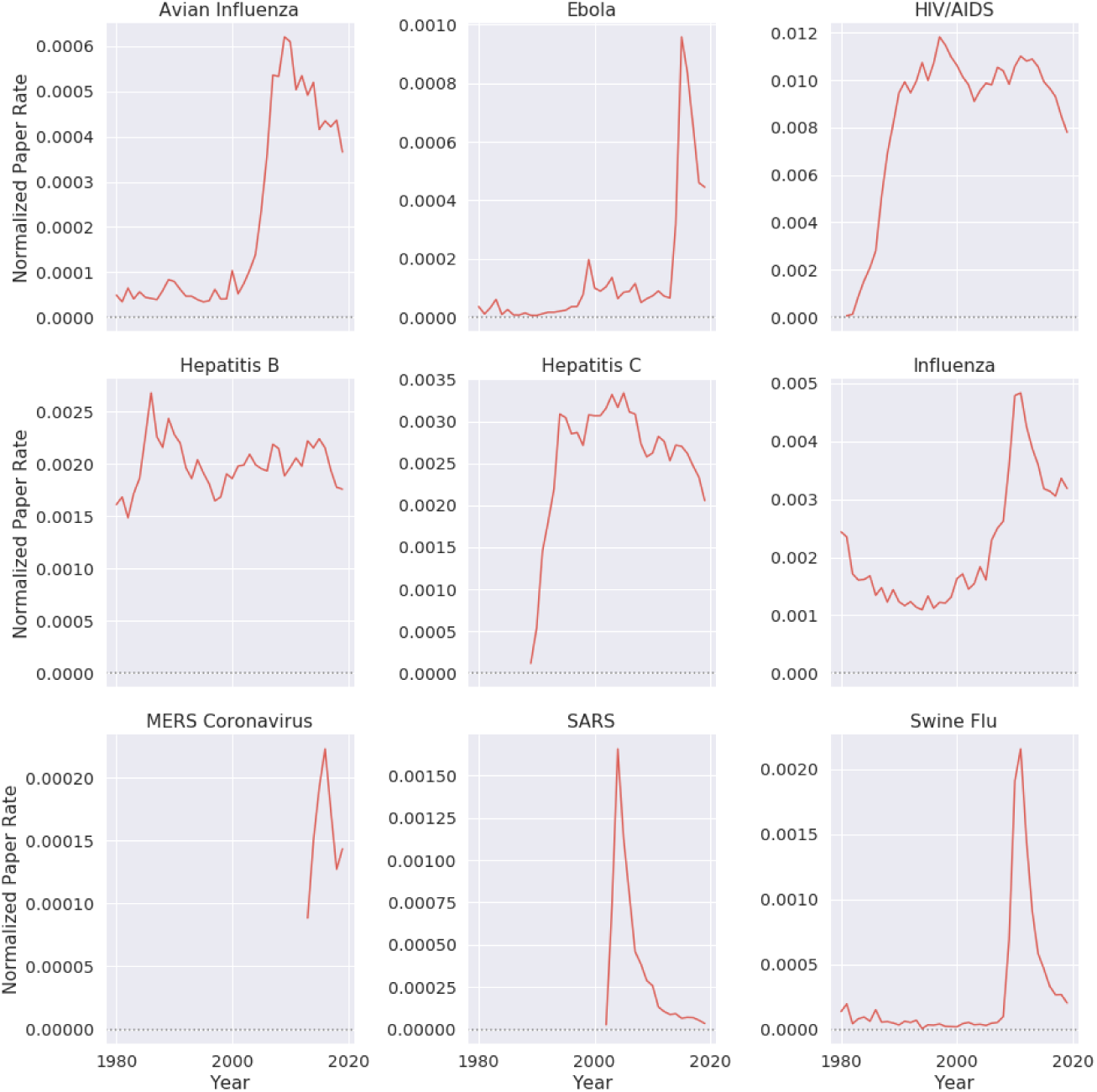
Normalized paper rate by different diseases over time. Diseases that have a drastic increase in their normalized number of publications mostly coincide with an epidemic.

**Figure 3:**
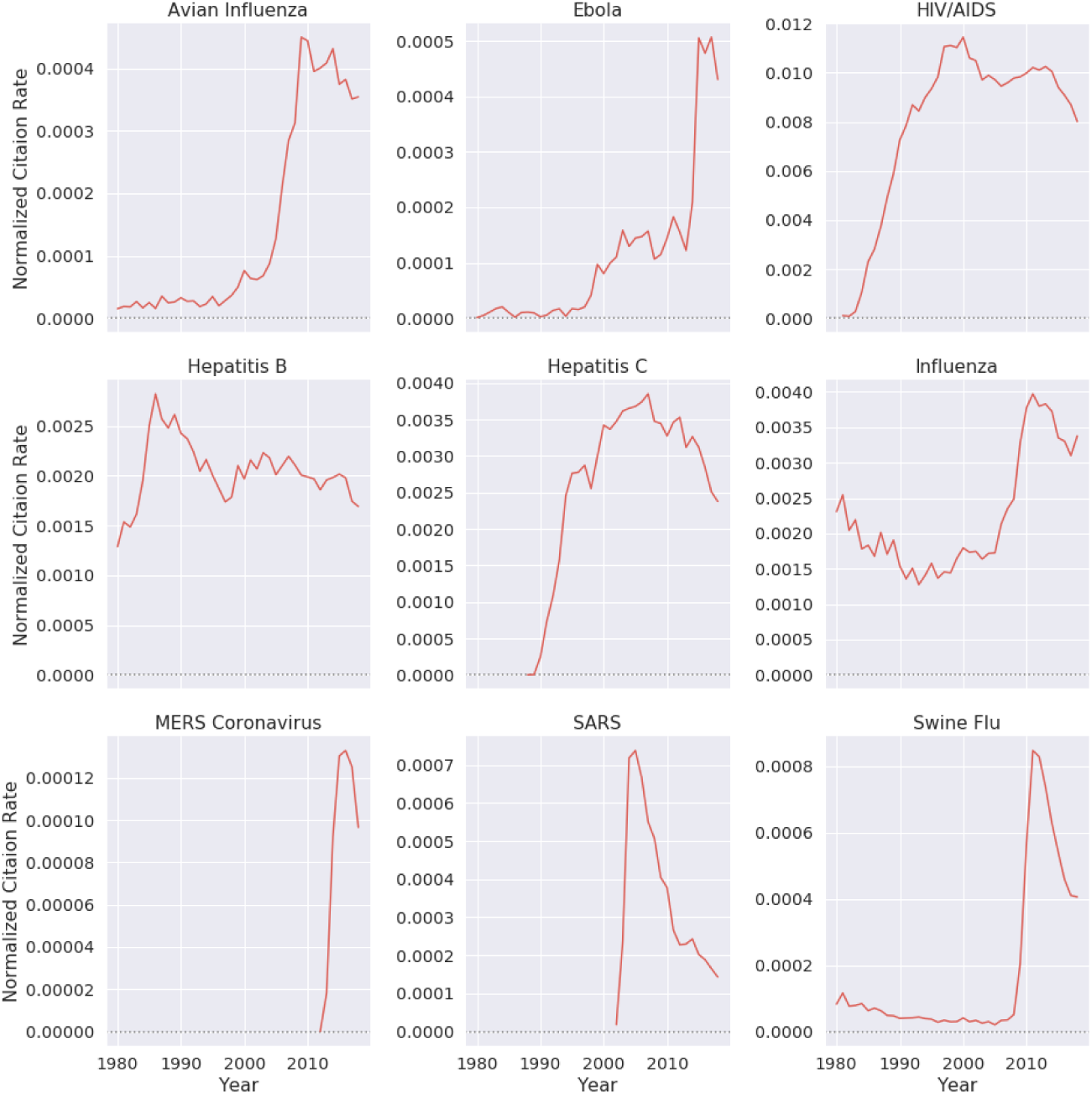
Normalized citation rate by different diseases over time. Diseases that have a drastic increase in their normalized number of citations mostly represent an outbreak.

**Figure 4:**
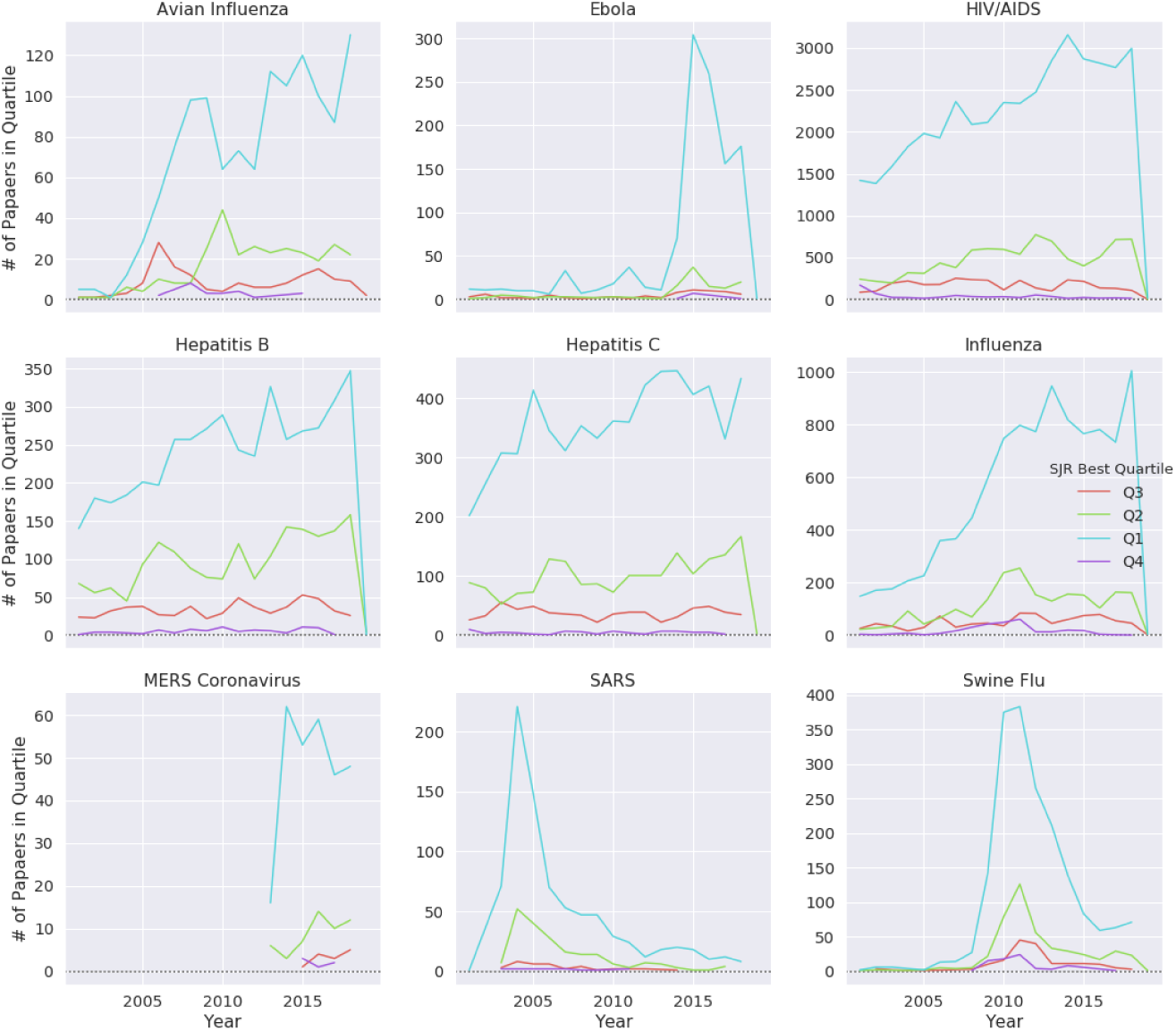
Publications by quartile over time for different diseases. Unlike other emerging infectious diseases, avian influenza did not demonstrate a decline in Q1 publications.

**Figure 5:**
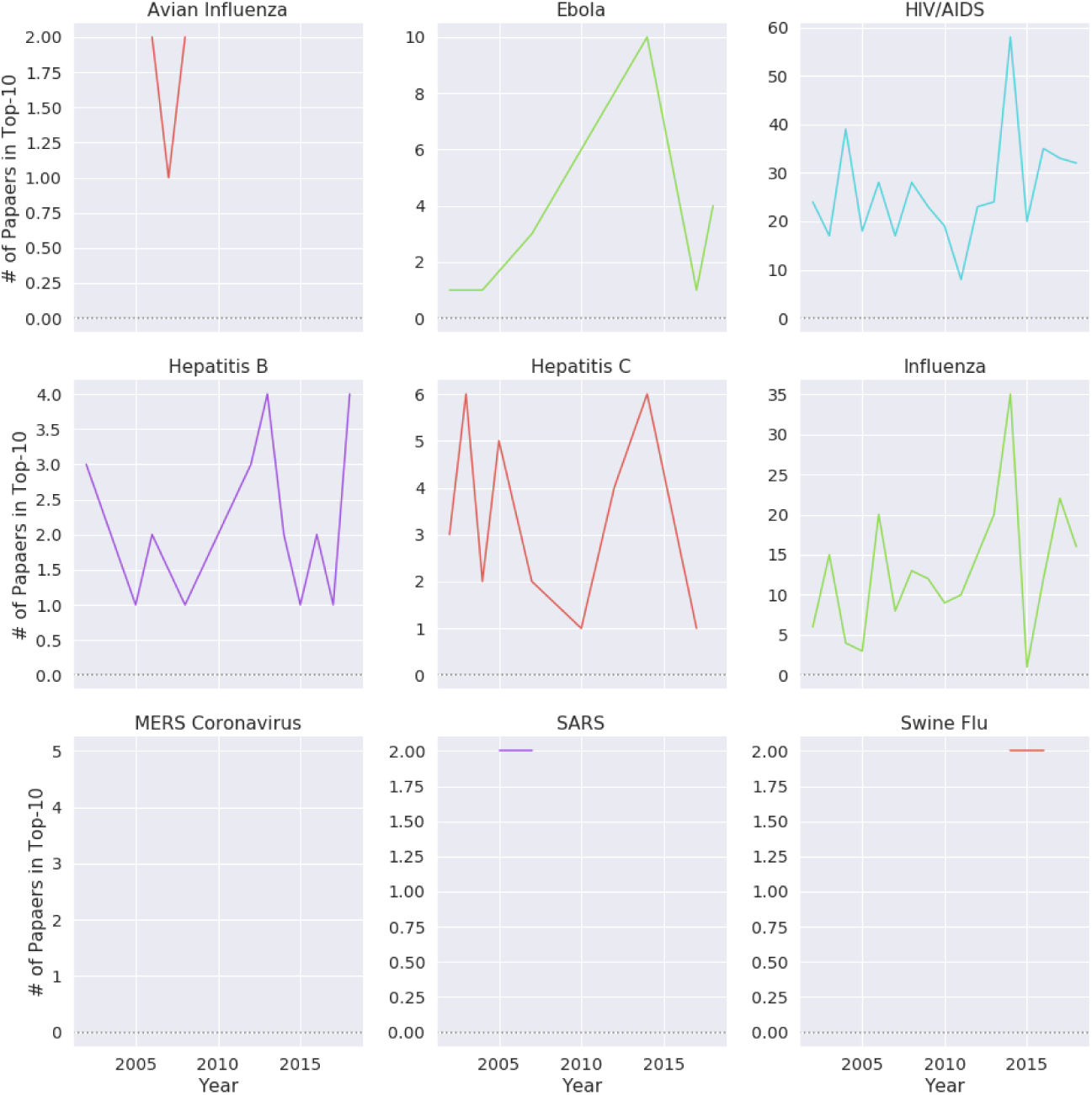
Number of papers by top-10 publications over time for different diseases.

**Figure 6:**
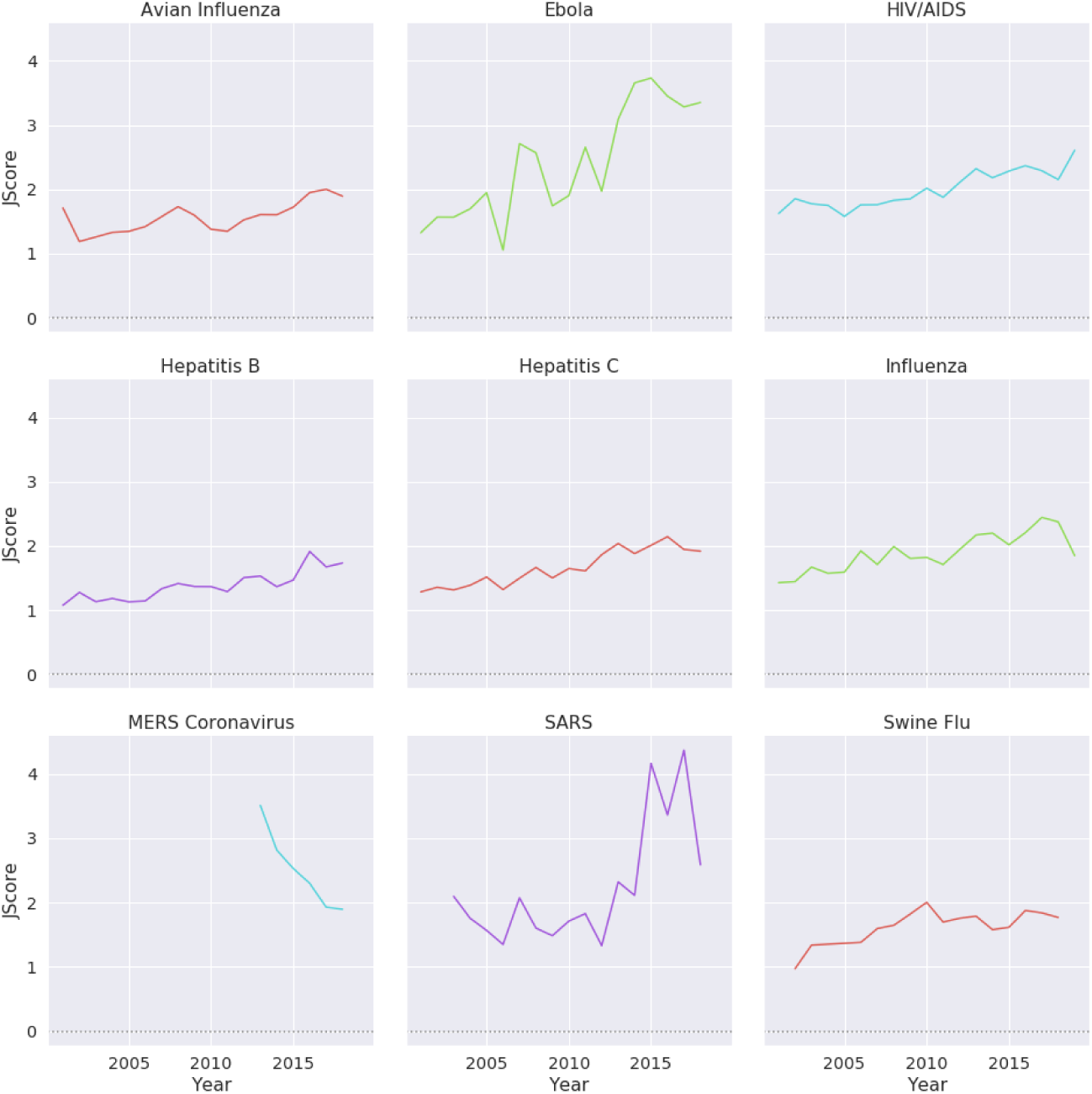
*JScore* over time for different diseases. Except for MERS, all presented diseases show an increase in JScore.

**Figure 7:**
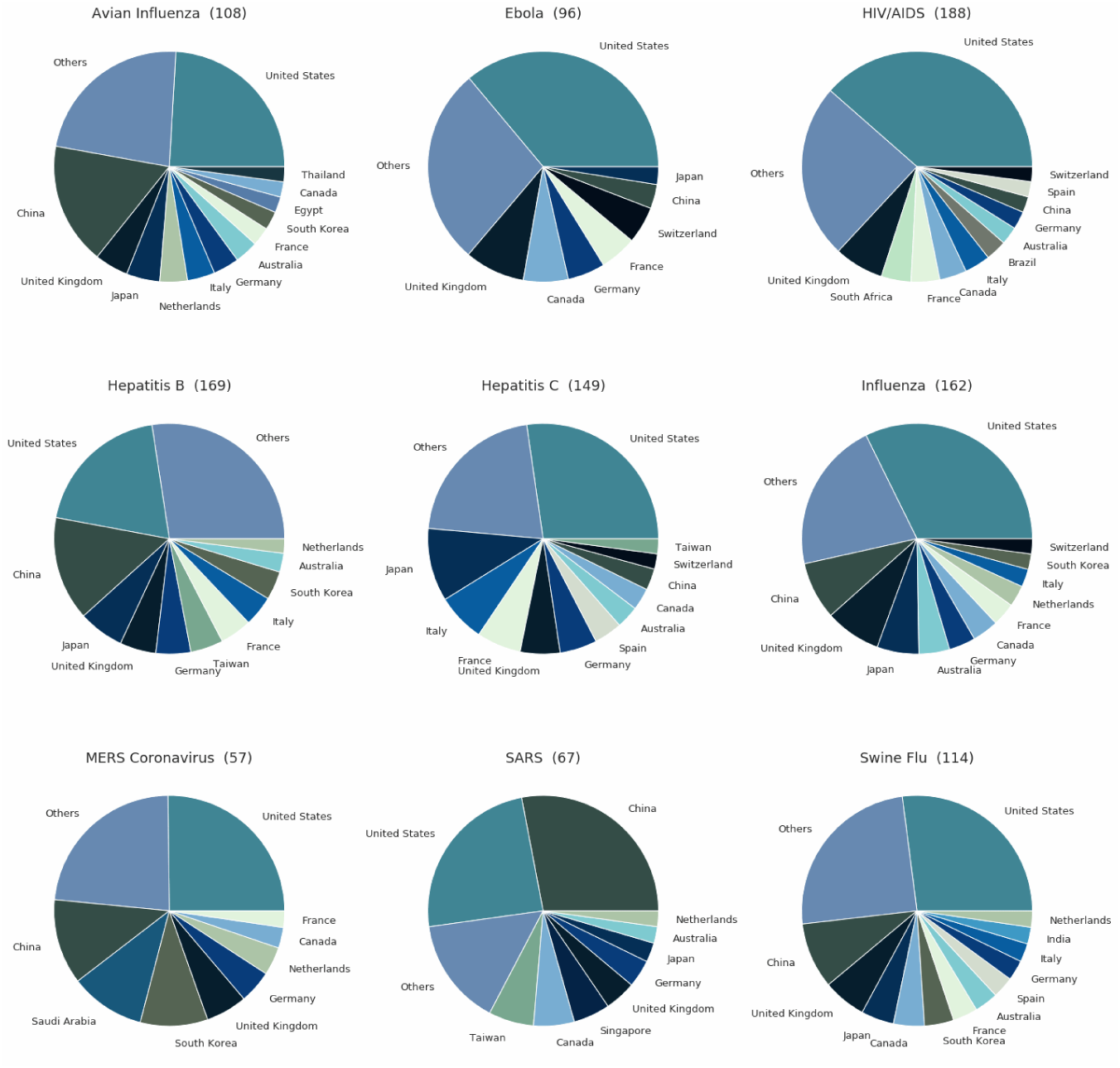
Number of researchers in each country for each disease. Most of the research was conducted in a small number of countries.

**Figure 8:**
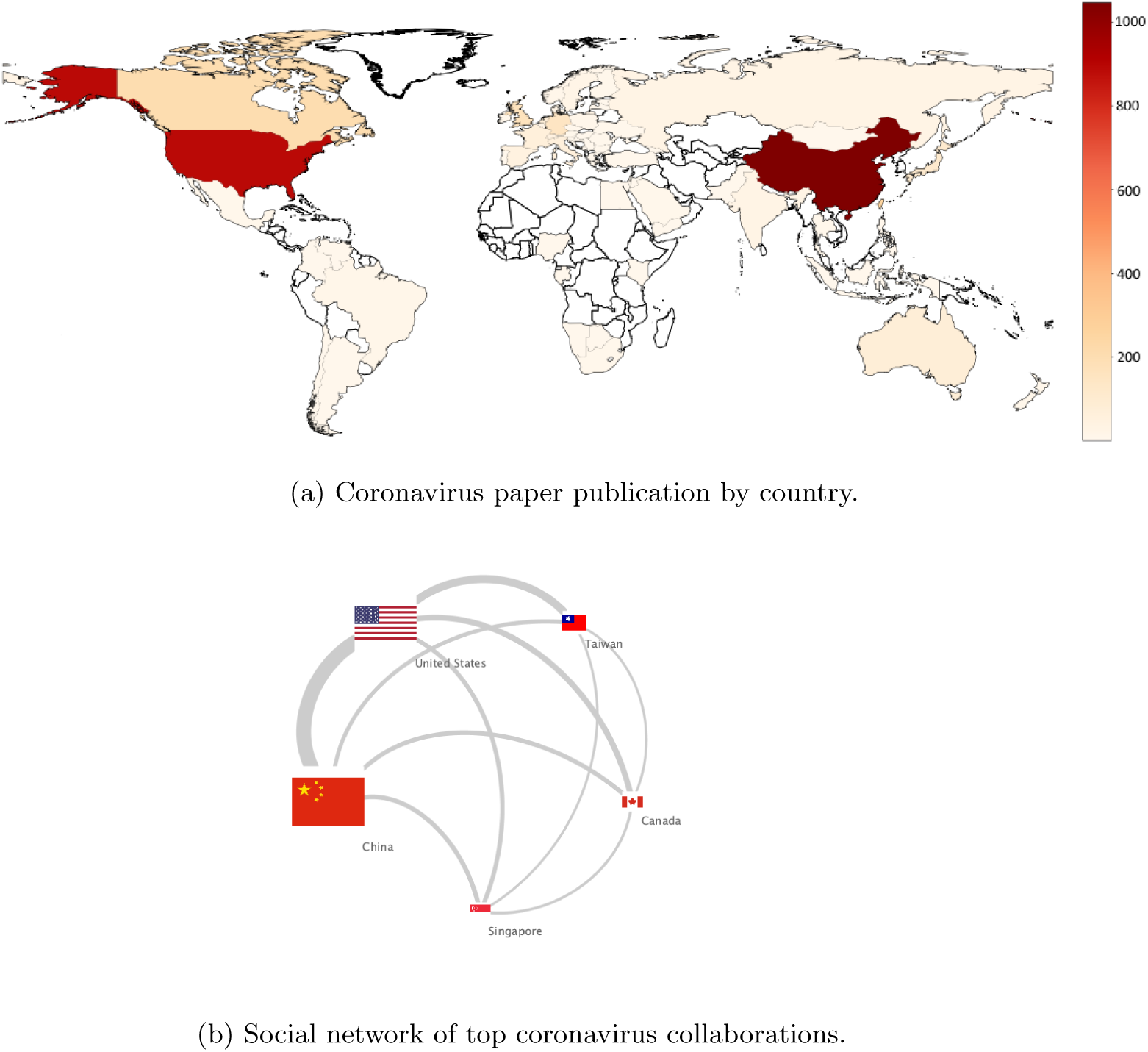
International research on the coronavirus.

### 5.2 Results of Journal Trends

From analyzing the trends in journal publications, we discovered the numbers of papers published by journal quartile are very similar to *Normalized Paper Rate* and *Normalized Citation Rate* (see Figure 4). We observed that for most of the diseases, the trends are quite similar: a growth in the study rate is coupled with a growth in the number of published papers in Q1 journals. We discovered that for SARS, MERS, the swine flu, and Ebola, Q1 publication trends were almost parallel to their NPR trends (see Figures 2 and 4). Also, we noticed that HIV, avian influenza, influenza, and hepatitis B and C have steady publication numbers in Q1 journals. Looking at papers in highly ranked journals (Figure 5), we observed that the diseases which are being continuously published in top-10 ranked journals are mainly persisting diseases, such as HIV and influenza. Additionally, we inspected how the average journal ranking of publications by disease has changed over time (Figure 6). We found that only MERS had a decline of *JScore*. We also noticed that current papers about SARS had the highest *JScore*.

### 5.3 Results of Author Trends

By studying the authorship trends in the research of emerging viruses, we discovered that there is a difference in the average experience of authors among diseases. SARS researchers had the lowest experience in years, and hepatitis C had the most experienced researchers (see Table 1). We noticed that the SARS research community had a smaller percentage of relatively prolific researchers than other diseases. Moreover, researchers with multiple papers related to SARS and MERS published on average 3.8 papers, while hepatitis C researchers published on average 5.2 papers during the same period. Additionally, from analyzing authors who published multiple papers on a specific disease, we found that on average there was a 2.5 paper difference between HIV and SARS authors. Furthermore, swine flu, SARS, and MERS were the diseases on which authors published the lowest number of multiple papers.

**Table 1:** Median researcher experience in years by disease.

| Disease | Median Experience in Years |
| --- | --- |
| SARS | 4 |
| Avian Influenza | 5 |
| Swine Flu | 5 |
| Hepatitis B | 5 |
| Ebola | 5 |
| Influenza | 6 |
| HIV/AIDS | 7 |
| MERS Coronavirus | 7 |
| Hepatitis C | 8 |

### 5.4 Results of Collaboration Trends

By inspecting global collaboration and research efforts, we found that the geolocation of researchers correlated with publication trends. For instance, most SARS, MERS, hepatitis B, and avian influenza research was done by investigators based in the US and China (Figure 7). In the case of SARS and MERS, most of the research stemmed from China and the US (Figure 8) with only about 17% of SARS papers’ first authors being located in Europe. Overall, researchers from 57 and 67 countries have studied MERS and SARS, respectively. However, the vast majority of SARS papers (73%) were written by researchers in only 6 countries (Figure 7). While the US was dominant in the research of all inspected diseases, China showed an increased output in only these three diseases. Also, MERS and SARS were studied in the least number of countries, and HIV was studied in the highest number of countries (Figure 7). Moreover, SARS and MERS were the diseases least studied in Europe, with only 17% and 19% of SARS and MERS studies, respectively, as opposed to Ebola studies, 29% of which were conducted in Europe.

## 6 Discussion

In this study, we analyzed trends in the research of emerging viruses over the past two decades with emphasis on emerging coronaviruses (SARS and MERS). We compared the research of these two coronavirus epidemics to seven other emerging viral infectious diseases as comparators. To this end, we used multiple bibliometric datasets, fusing them to get additional insights. Using this data, we explored the research of epidemiology from the perspectives of papers, journals, authors, and international collaborations.

By analyzing the results presented in the Results section, the following can be noted: First, the surge in infectious disease publications (Figure 1) supports the results of Fire and Guestrin [36] that found there has been a general escalation of scientific publications. We found that the growth in the number of infectious disease publications is very similar to other fields. Hence, Goodhart’s Law^6^ did not skip the world of virology research. However, alongside the general growth in the number of papers, we observed that there was a decline in the relative number of papers on the specific infectious diseases we inspected. The most evident drastic drop in the publication rate happened after an epidemic ended. It appears that, for a short while, many researchers study an outbreak, but later their efforts are reduced. This is strengthened by considering the average number of multiple papers per author for each disease (see Table 2). Additionally, similar patterns were found in the funding of MERS and SARS research [25], which indicates that there is a possibility that the research rate has decreased due to lack of funding.

**Table 2:** Average papers published by author with multiple papers related to a specific disease.

| Disease | Papers |
| --- | --- |
| Swine Flu | 3.45 |
| SARS | 3.84 |
| MERS Coronavirus | 3.86 |
| Ebola | 4.07 |
| Hepatitis B | 4.42 |
| Avian Influenza | 4.47 |
| Influenza | 5.04 |
| Hepatitis C | 5.24 |
| HIV/AIDS | 6.31 |

Second, when looking at journal publications, we noted very similar patterns occurred for citations and publications. This result emphasizes that fewer publications, and hence fewer citations, translate into fewer papers in Q1 journals (Figure 4). Also, we observed the same patterns as Fire and Guestrin [36], with most of the papers being published in Q1 journals and the minority published in Q2-Q4 journals. This trend started to change when zooming in and analyzing publications in top-10 ranked journals (Figure 5). While we can see some correlation to outbreaks in Ebola, swine flu, and SARS, it is harder to interpret the curve of HIV since there were no focused epidemics in the past 20 years but a global burden, and we did not observe similar patterns in publications and citations. Observing the *JScore* (Journal Trends Section) results (Figure 6), most diseases showed a steady increase, but two diseases behaved rather anomalously. MERS had a decline since 2013, which is reasonable to expect after the initial outbreak, but we did not see the same trend in the other diseases and there is a general trend of increasing average SJR [36]. The second anomaly is that SARS had an increase in *JScore* alongside a decrease in citations and publication numbers. Inspecting the data, we discovered that in 2017 there were three published papers in Lancet Infectious Diseases and in 2015 two papers in Journal of Experimental Medicine about SARS, and both journals have a very high SJR. These publications increased the *JScore* drastically. This anomaly is a result of outliers in the data that biased the results. We can observe in Figure 4 that in the last decade the number of SARS papers published in ranked journals dropped drastically. It dropped low enough that two outliers created a bias on the *JScore*. Generally, the less data we have, the greater chance for outliers to cause bias in the data.

Third, we observed that on average authors write a fewer number of multiple papers on diseases that are characterized by large epidemics, such as the swine flu and SARS. On the other side of the scale are hepatitis C and HIV, which are persistent viral diseases with high global burdens. These diseases involve more prolific authors. Regarding Ebola and MERS, it is too early to predict if they will behave similarly to SARS since they are relatively new and require further follow up. Fourth, looking at international collaboration, we observed the US to be very dominant in all the disease studies (Figure 7). Looking at China, we found it to be mainly dominant in diseases that were epidemiologically relevant to public health in China, such as SARS, avian influenza, and hepatitis B. When looking at Ebola, which has not been a threat to China for the last two decades, we observed a relatively low investment in its research in China. We observed that regarding MERS, we found similar results to Sa’ed [50]. In both studies the top-3 biggest contributors in MERS studies were the US, China, and Saudi Arabia.

Many of the trends we observed are related to the pattern of the diseases. We observed two main types of infectious diseases with distinct trends. The first type was emerging viral infections like SARS and Ebola. Their academic outputs tend to peak after an epidemic and then subside. The second type were viral infections with high burdens such hepatitis B and HIV, for which there is a more or less constant trend. These trends were most evident in publication and citation numbers, as well as journal metrics. The collaboration and author distributions were more affected by where the outbreak occurred or where there was a high burden. This was also strengthened in the clusters we found where they were divided in the same way.

In terms of practical implications, we see several options. First, notwithstanding the importance of pathogen discovery, as evident in projects like the Global Virome Project [51] that is trying to discover unknown zoonotic viruses to stop future outbreaks, it is still important to monitor the status of current research that concerns known pathogens. It can be observed from Figures 2 and 3 that there are diseases with declining interest from the scientific community. These trends are harder to spot when looking at the total number of publications since the total number of papers generally keeps growing (Figure 1a). Using NPR and NCR can help decision makers investigate if additional resources should be invested in the study of these diseases. For instance, while SARS and MERS were in WHO’s R&D Blueprint as priority diseases, they still exhibited a decline in their research rate. Second, using collaboration data, it is possible to find which countries have potential for growth in the number of researchers on specific diseases and also which bilateral grants have potential.

Currently, there is no doubt that we have to be better prepared for the next pandemic and the emergence of “Disease X.” We observed that currently there is a non-sustained investment in EIDs such as SARS and MERS, which is a key issue. Another crucial issue is the sharing of research material such as data and code. Data and code allow scientists to make more accurate discoveries faster by continuing knowledge from previous studies. Using the MAG dataset Paper Resources table, we inspected how many papers from the nine diseases we analyzed had code or data. We found that there were 30 and 75 papers that had data and code, respectively. These numbers are very low, and we suspect that there are a lot of missing data in this table. We firmly believe that publishing code and data should be mandatory when possible.

This study may have several limitations. To analyze the data, we relied on titles to associate papers with diseases. While a title is very important in classifying the topic of a paper, some papers may discuss a disease without mentioning its name in the title. Additionally, there may be false positives; for instance, an acronym might have several meanings that are not related to an infectious disease term. An additional limitation is our focus on a limited number of distinct diseases. There are other emerging infections not evaluated here in which could have followed other trends. To deal with some of these limitations, we only analyzed papers that were categorized as medicine and biology papers as a means to reduce false positives. Furthermore, we show that the same trends appeared even when we filtered all the papers by the category of virology (see Figures 11 and 12). Finally, we compared papers that were tagged with a MeSH term on PubMed to the papers we retrieved using our keyword search of the title. We found that we matched MeSH terms with 73% recall, which is in the range described by Breugelmans et al. [24].

In the future, we would like to perform extended collaboration analysis by improving the institution country mapping. Currently, we were able to identify 94.2% of the countries of origin for the institutions in the MAG affiliation table. We intend to improve the institution country mapping by using additional data sources. Additionally, we are planning to extend our study into other diseases and look for correlations with real-world data such as global disease burden.

## 7 Conclusions

The COVID-19 outbreak has emphasized the insufficient knowledge available on emerging coronaviruses. Here, we explored how previous coronavirus outbreaks and other emerging viral epidemics have been studied over the last two decades. From inspecting the research outputs in this field from several different angles, we demonstrate that the interest of the research community in an emerging infection is temporarily associated with the dynamics of the incident and that a drastic drop of interest is evident after the initial epidemic subsides. This translates into limited collaborations and a non-sustained investment in research on coronaviruses. Such a short-lived investment also involves reduced funding as presented by Head et al. [25] and may slow down important developments such as new drugs, vaccines, or preventive strategies. There has been an unprecedented explosion of publications on COVID-19 since January 2020 and also a significant allocation of research funding. We believe the lessons learned from the scientometrics of previous epidemics argue that regardless of the outcome of COVID-19, efforts to sustain research in this field should be made. More specifically, in 2017 [52] and 2018 [53], SARS and MERS were considered to be priority diseases in WHO’s R&D Blueprint, but their research rate did not grow relative to other diseases. Therefore, the translation of international policy and public health priorities into a research agenda should be continuously monitored and enhanced.

## 8 Data Availability

The datasets supporting the results of this article are available online (see Data Description section). Preprocessed datasets is a available on [54].

## 9 Availability of source code and requirements

- Project name: ScienceDynamics
- Project home page: https://github.com/data4goodlab/ScienceDynamics
- Operating system(s): Linux, OS X
- Programming language: Python
- Other requirements: Python 3.6 or higher
- License: MIT License
- RRID:SCR 018819

## 10 Acknowledgements

We would like to thank Carol Teegarden for editing and proofreading this article to completion and to David Schmid for sharing with us the MAG dataset.

## A Appendix

**Figure 9:**
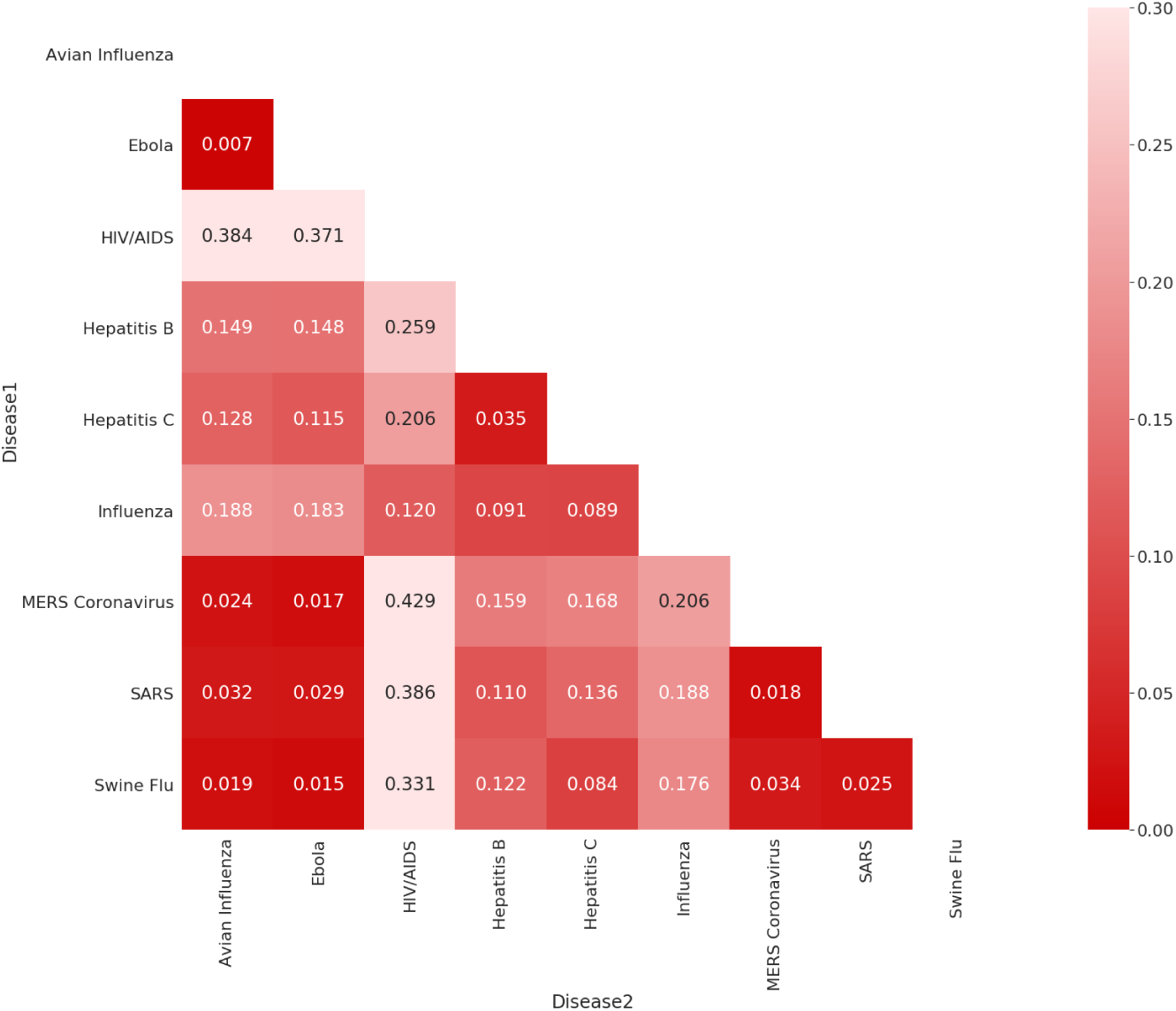
DTW distance between NPR of diseases

**Figure 10:**
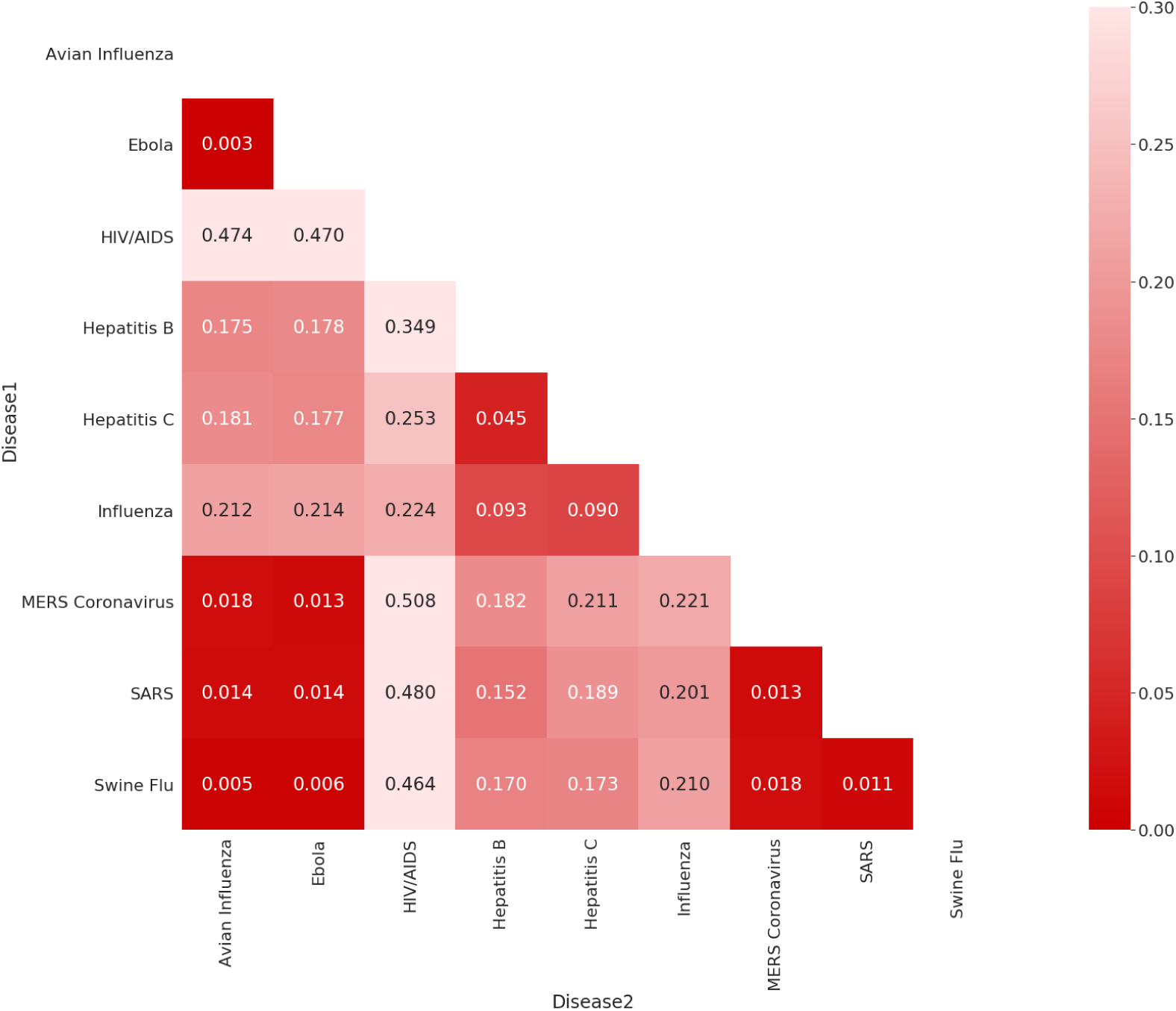
DTW distance between NCR of diseases.

**Figure 11:**
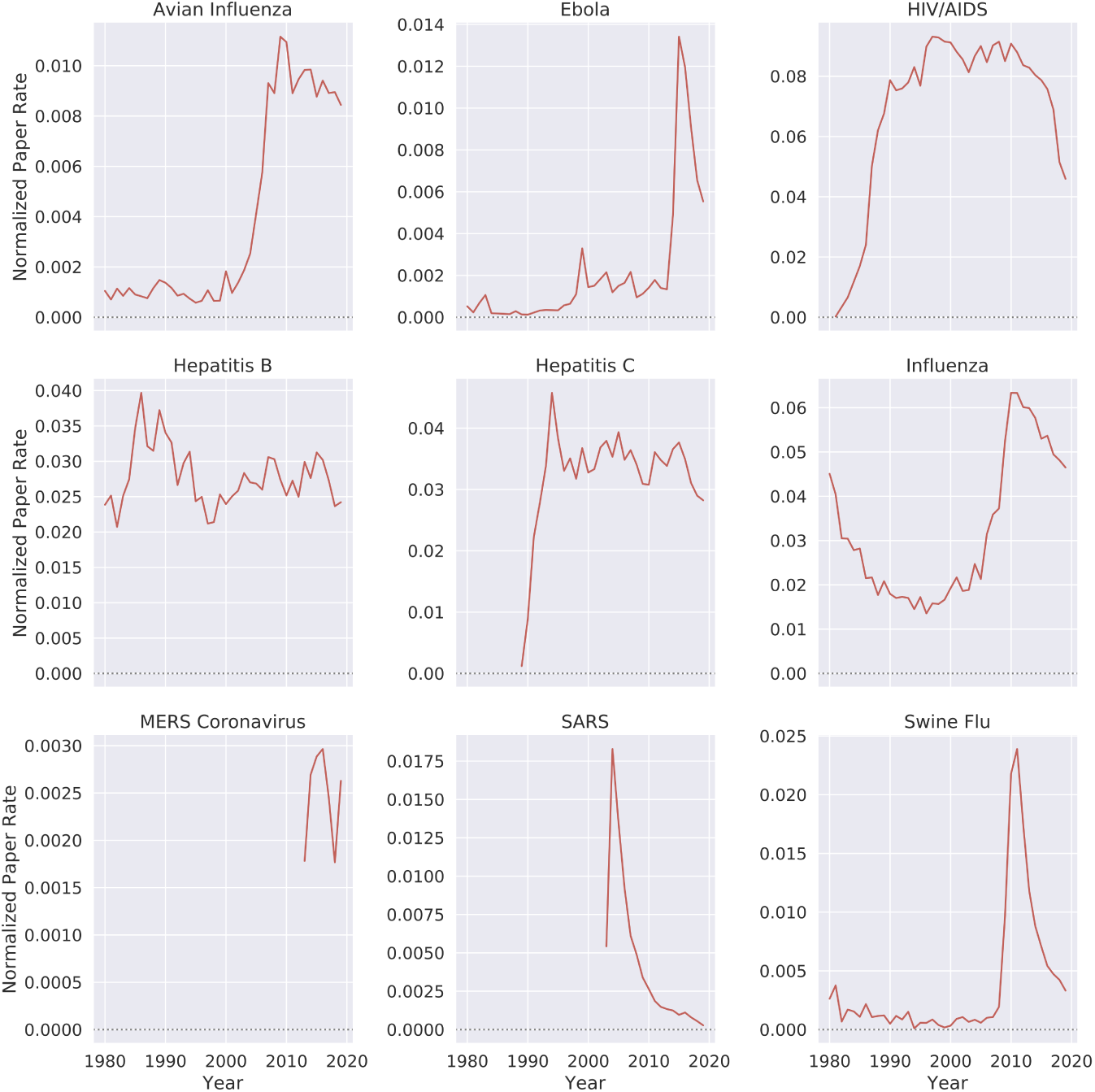
Normalized paper rate of the virology category by different diseases over time.

**Figure 12:**
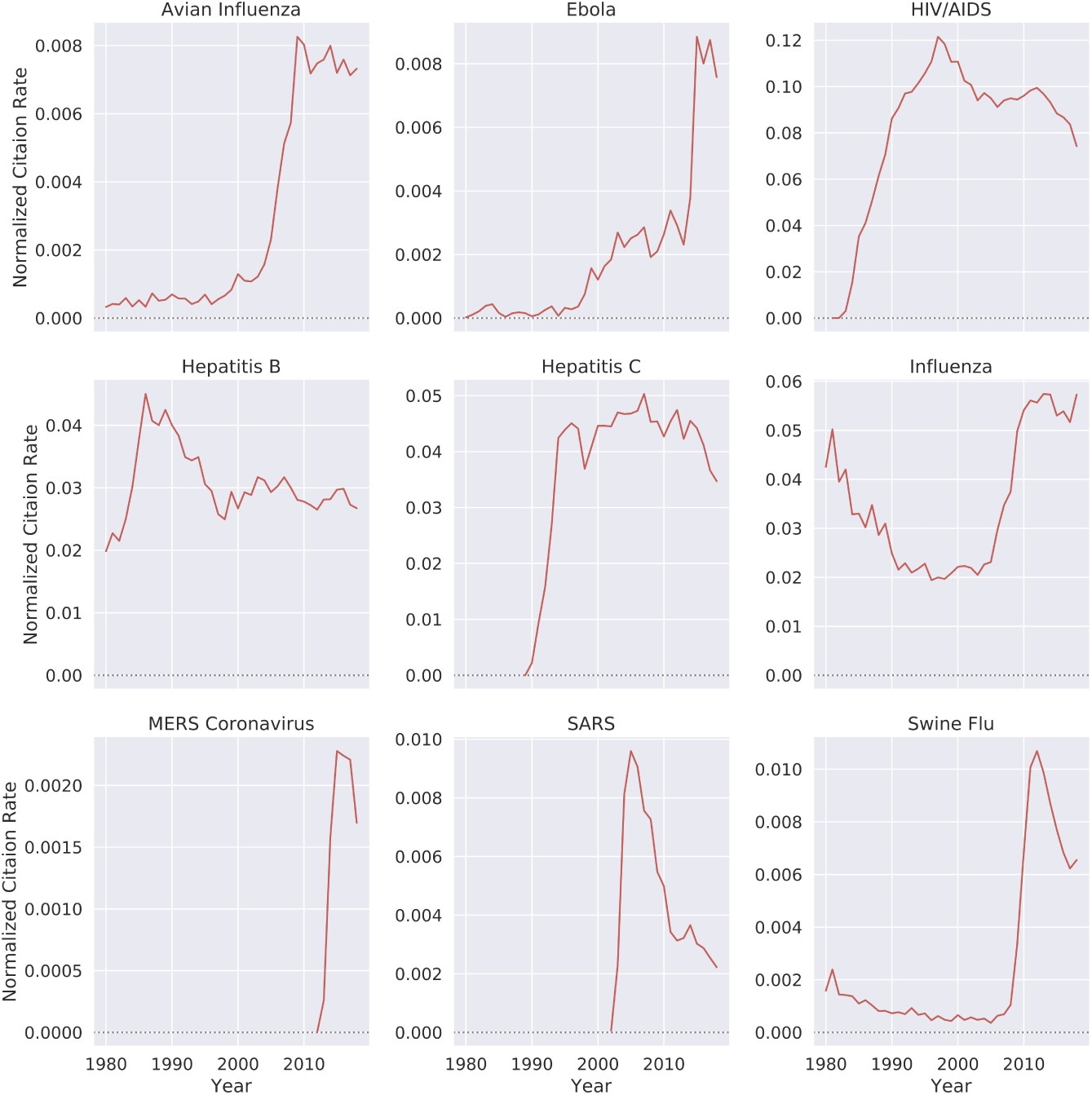
Normalized citation rate of the virology category by different diseases over time.

## Footnotes

1 The SJR indicator is a measure used to assess the prestige of a journal. The measure takes into account the number of citations and the prestige of the source of the citing paper [42]

2 ”The Journal Impact Factor quartile is the quotient of a journal’s rank in category (X) and the total number of journals in the category (Y), so that (X / Y) = Percentile Rank Z” [43].

3 To determine which papers, we used the MAG fields of study.

4 The fields used were “coordinate location (P625),” “country (P17),” “located at street address (P6375),” “located in the administrative territorial entity (P131),” “headquarters location (P159),” and “location (P276).”

5 English Wikipedia

6 ”When a measure becomes a target, it ceases to be a good measure.”

